# Linking genotype to longevity under genealogical discordance in *Sebastes* rockfishes

**DOI:** 10.64898/2026.03.26.714443

**Authors:** Yu K. Mo, Peter H. Sudmant, Matthew W. Hahn

## Abstract

Rockfishes (genus *Sebastes*) show extreme variation in longevity among closely related species, but the evolutionary history of this young radiation is highly complex. To unpack these relationships and to associate genotypes with phenotypes, we quantified genealogical discordance among 55 *Sebastes* species and implemented a phyloGWAS framework that incorporates discordant gene histories into genotype–longevity association tests. We found that genealogical discordance is extremely high: the inferred species tree topology differed among several ILS-aware methods, with most internal branches having low concordance factors regardless of which method was used. Nevertheless, some phylogenetic structure was shared by all inferred species trees. We used simulations to assess the statistical properties of phyloGWAS applied to complex traits using different genetic relatedness matrices (GRMs) and under varying levels of discordance. Adding an accurate GRM reduced false positives relative to a model without relatedness, but GRMs only modestly increased power to detect true positives. Using multiple approaches on the *Sebastes* data, phyloGWAS identified several variants associated with longevity. Our results indicate that extreme genealogical discordance is a core feature of *Sebastes* evolution and that phyloGWAS can help in connecting genotype to phenotype under these conditions.

## Introduction

Rockfishes in the genus *Sebastes* are an exceptional system for studying the evolution of life-history traits. Despite being a relatively recent radiation of approximately 100 species in the northern Pacific Ocean, lineages span more than an order of magnitude in maximum lifespan, from 10 years to 200 years (Love et al. 2002; Mangel et al. 2007). As a result of this remarkable diversity in lifespan, rockfishes have become a model system for this life-history trait and its determinants. In order to understand both the genetic and ecological factors underlying lifespan, a clear understanding of the phylogenetic relationships among the rockfishes is necessary (cf. Felsenstein 1985).

The rockfish radiation is estimated to have begun only 8-10 million years ago (Hyde and Vetter 2007; Kolora et al. 2021), though some estimates place the base of the radiation much more recently (Johns and Avise 1998). Regardless of the exact timing, the large number of species and speciation events that followed means that the time between speciation events will be very short. Given the long generation times and moderate effective population sizes of rockfishes (Gomez-Uchida and Banks 2006), these short internode distances will be exacerbated, resulting in a large amount of incomplete lineage sorting (ILS) and gene tree-species tree discordance among loci (Doyle 1992; Maddison 1997; Degnan and Rosenberg 2009). Although previous studies of *Sebastes* have used multiple genes or markers to construct a species tree (Kolora et al. 2021; Wallace et al. 2022; Treaster et al. 2023), none has quantified the amount of genealogical discordance in this clade.

Estimating and accommodating gene tree discordance is important in phylogenetics, especially when trying to understand the evolutionary history of genes and traits (like lifespan). If one uses only a single, fixed species tree in comparative analyses, then all genes and traits are forced to follow the species tree (Hahn and Nakhleh 2016). However, when many gene trees do not match the species tree, then standard phylogenetic methods that use only a single tree can be misled about the number of times a trait has evolved, the direction of evolution, the rate of evolution, and the timing of trait transitions (Guerrero and Hahn 2018; Mendes et al. 2018; Hibbins et al. 2020, 2023; Adams et al. 2025; Schraiber et al. 2024; Mo and Hahn 2025). For instance, there are now multiple examples of ancestrally polymorphic traits that are differentially fixed among descendant lineages (e.g. Fontaine et al. 2015; Lamichhaney et al. 2016; Li et al. 2016; Pease et al. 2016; Han et al. 2017; Palesch et al. 2018; Wu et al. 2018; Louis et al. 2021; Feng et al. 2022). Using only a single species tree and ignoring gene tree discordance would incorrectly lead to the conclusion that such traits had arisen convergently multiple times.

Here, we investigate species relationships and genealogical discordance in *Sebastes*. We construct species trees for *Sebastes* with three different methods—all of which are intended to accurately infer a species tree in the presence of ILS—and estimate the amount of genealogical discordance across the clade. In addition to simply quantifying discordance, we take advantage of this gene tree heterogeneity to search for genetic variants associated with lifespan: we implement a phyloGWAS framework (Pease et al. 2016; Wu et al. 2018) to test genotype–phenotype associations for longevity while explicitly incorporating discordant relationships. We demonstrate via simulations the performance of genotype–phenotype association models under varying genealogical discordant scenarios and using various approaches to control for discordance. Application of these approaches to the *Sebastes* data revealed several promising candidate variants associated with longevity.

## Results

### Four major groups across 55 Sebastes species

Across 55 *Sebastes* species and the outgroup *Sebastolobus alascanus* (all from Kolora et al. 2021; Materials and Methods), we were able to analyze 9706 orthologous genes with at least 50% of the alignment covered in 70% of species. Among variable sites, 33.1% were parsimony-informative. A concatenated alignment of all these data was used as input to CASTER-site (Zhang et al. 2025) and SVDquartets (Chifman and Kubatko 2014). Gene trees were inferred separately for each locus using IQ-TREE 2 (Minh et al. 2020a), and were used as input to ASTRAL (Mirarab et al. 2014).

The three methods used for inferring species trees gave three different tree topologies, with both shared and conflicting structures (Figure 1; Supplementary Figure 1). Arbitrarily choosing the CASTER tree as a reference, we can see the fairly consistent presence of four monophyletic groups (Figure 1a). The yellow group (17 species) retains a largely identical backbone across methods; nevertheless, it represents species placed in at least three different subgenera (Hyde and Vetter 2007). The green group (17 species) preserves a consistent structure, at least within shallow subclades. The purple group (10 species) matches between SVDquartets and CASTER, but becomes paraphyletic under ASTRAL. Within the blue group (11 species), the subclade containing *S. melanops, S. flavidus, S. mystinus*, and *S. entomelas* matches across methods.

**Figure 1.**
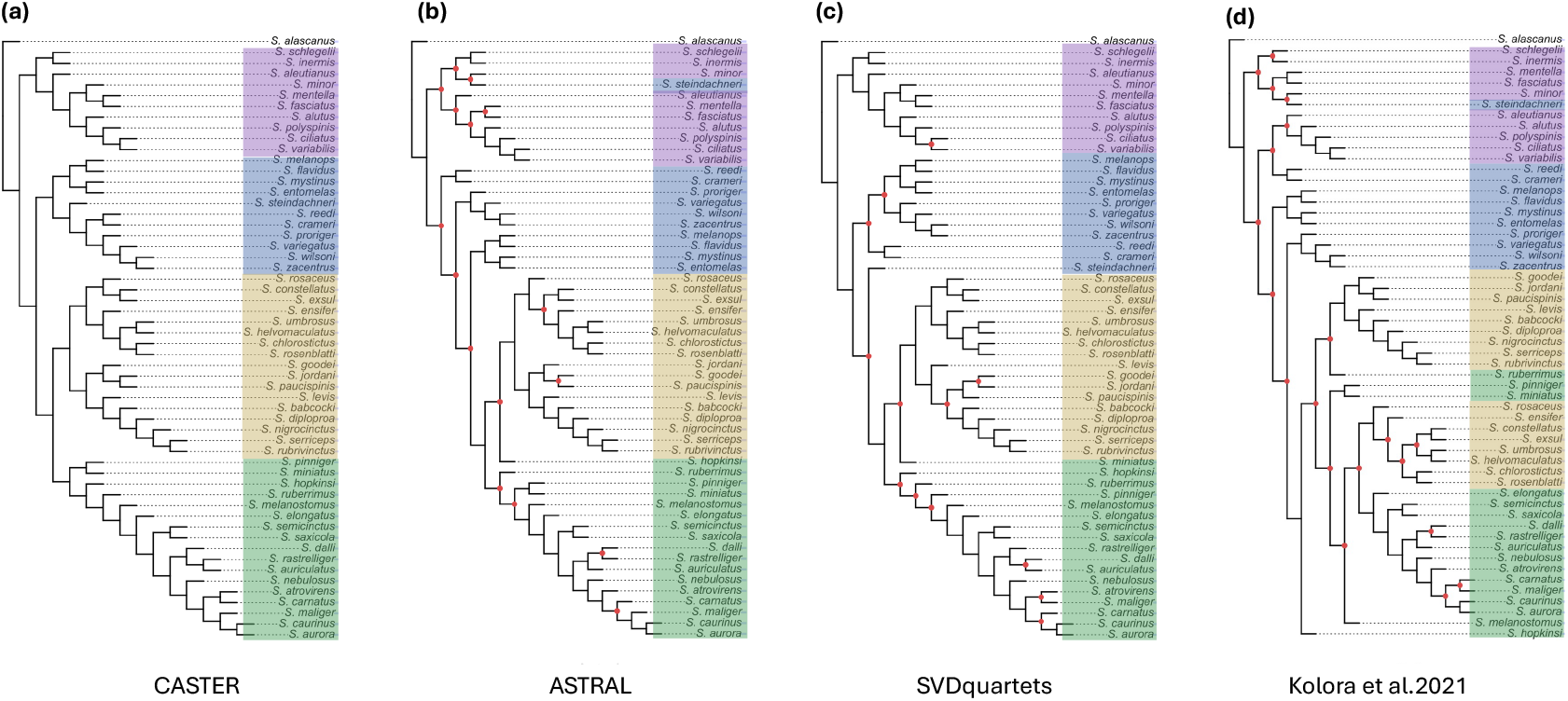
Species trees inferred by different methods. Species trees among 55 *Sebastes* species inferred from 9706 orthologous genes using (a) CASTER, (b) ASTRAL, and (c) SVDquartets; all trees are rooted with the outgroup *Sebastolobus alascanus*. Colored clades follow the monophyletic groups defined in (a). (d) Species tree reported in Kolora et al. (2021). (b)-(d) Red circles mark nodes that do not exist in (a).

Several substructures conflict across methods (Figure 1; Supplementary Figure 1). In the yellow group, the clade containing *S. goodei, S. jordani*, and *S. paucispinis* shows all three alternative topologies. In addition, the position of *S. levis* conflicts when using SVDquartets. In the green group, the placement of *S. hopkinsi, S. miniatus*, and *S. pinniger* differs among methods; SVDquartets also places *S. dalli* as sister to *S. auriculatus* rather than *S. rastrelliger*, and conflicts with CASTER by placing *S. mailiger* closer to *S. aurora* and *S. caurinus*. In the purple group, the relative paraphyly in ASTRAL stems from the placement of *S. steindachneri* (assigned to the blue group in CASTER), while also altering several other minor clades. The blue group is no longer monophyletic at all under ASTRAL, while SVDquartets has again a rearrangement of *S. steindachneri* relative to CASTER (Supplementary Figure 1a,b).

We also compared our results with the species tree from Kolora et al. (2021), which also used ASTRAL to reconstruct the tree. There are many rearrangements between this tree and all three trees inferred here (Figure 1d; Supplementary Figure 1c), though note that in both cases ASTRAL has placed *S. steindachneri* in the purple clade. Although the data we use here originate from the study of Kolora et al., the tree presented in their paper used only 670 orthologs identified using the BUSCO pipeline (Waterhouse et al. 2018). Given that we have more than 10 times as many gene trees, we think that the trees presented here may be a more accurate representation of the species tree. However, given the very high discordance we find (see next section), the exact branching pattern in the species tree may not be very predictive of the evolution of traits, regardless of which tree is the correct one (cf. Lanfear and Hahn 2024).

### High discordance across rockfishes

Given the rapid radiation of a large number of rockfish species in a short time period, we expect to see a large amount of genealogical discordance. Indeed, there was extensive topological conflict in the *Sebastes* species tree (Figure 2). We first quantified site concordance factors (sCFs) on the CASTER tree (Figure 2a). sCFs measure the fraction of informative nucleotide sites for a single branch that support the species tree, while the two site discordance factors (sDF1 and sDF2) represent the fraction of sites supporting either of the two alternative topologies that are one nearest-neighbor interchange away for the same branch (Minh et al. 2020b; Mo et al. 2023). Across all branches of the *Sebastes* tree, sCFs are very low (median 36.1%, IQR 16.3%; mean 40.2%; Supplementary Figure 2a). Over one-third of internal branches (18 of 53) fall below 33.3%, and only one branch exceeds 75% (Supplementary Figure 2a). For reference, a completely unresolved star tree should have an sCF of 33.3%; values below this indicate that the topological resolution present in the species tree is not the single most common resolution among individually informative sites.

**Figure 2.**
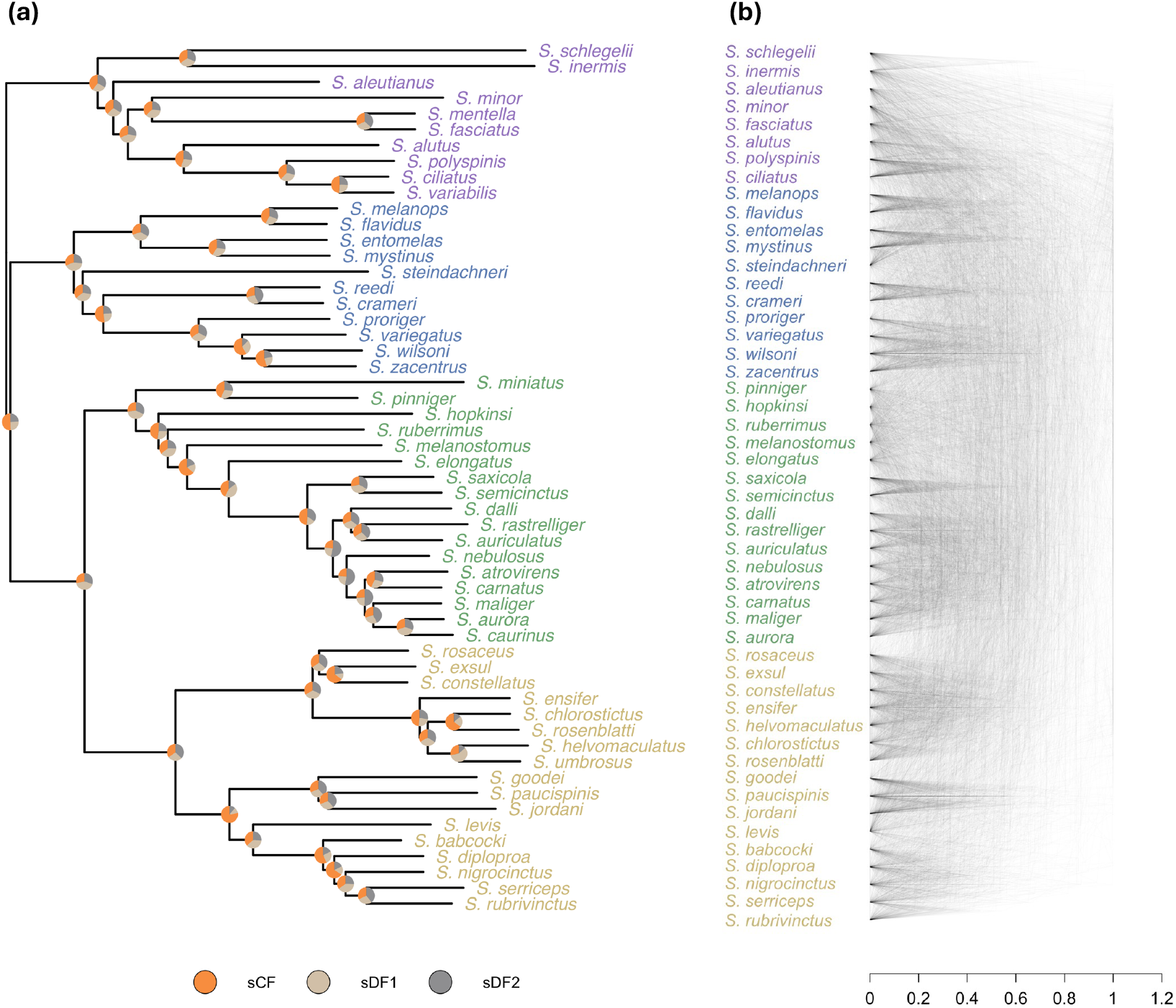
Discordance across the species tree. (a) CASTER species tree with branch lengths in substitutions per site estimated by IQ-TREE 2. Pie charts at internal nodes show site concordance factors (orange) and site discordance factors for the two minor topologies (grey). (b) Cloudogram summarizing 389 gene trees for 50 *Sebastes* species with full coverage across species.

If we use any of the other possible species tree topologies (ASTRAL, SVDquartets, or Kolora et al.), we get comparable sCF values (Supplementary Figure 2a). The mean and median sCF across branches are largely the same using these trees as the species tree, though often there are a few more branches with higher sCFs and many more branches with lower sCFs. These results preclude a straightforward determination of the “best” tree as the one with the highest average sCF, though perhaps the absence of very low sCFs from the CASTER tree provides a bit of support in its favor.

We also calculated gene concordance factors (gCFs), which represent the fraction of informative gene trees that match the species tree for each branch (Ané et al. 2007). For the CASTER tree, gCFs are very low (median 20.1%, IQR 23.1%; mean 13.9%; Supplementary Figure 2b), with nearly identical distributions across methods (Supplementary Figure 2b). Because there are many possible topologies in a gene tree, even highly supported branches in a species tree can have gCFs that fall below 33.3% (Lanfear and Hahn 2024). However, we also find substantial proportions of branches in the species tree inferred by each method that have less than 5% of gene trees supporting them: 24.1% of internal branches for the CASTER tree, 22.2% for the ASTRAL tree, 25.9% for the SVDquartets tree and 24.1% for the tree in Kolora et al. 2021. This is not expected, though mathematically possible in extremely rapid radiations (e.g. Jarvis et al. 2014). As a visual representation of this extreme discordance, Figure 2b shows a “cloudogram” (Maddison 1997) among several hundred gene trees in *Sebastes*. While some structure is evident in shallow clades, it very quickly becomes difficult to discern any consistent phylogenetic histories.

### Genotype–phenotype associations with phyloGWAS

Longevity varies widely within the *Sebastes* groups, ranging from 12 to 205 years (Figure 3). A few attempts have been made to understand the ecological and genetic basis for this variation in longevity using phylogenetic comparative methods, but in each case a single species tree has been assumed to be the correct model for trait evolution (Mangel et al. 2007; Kolora et al. 2021; Treaster et al. 2023). However, the results above suggest that this trait (or the sub-traits that together determine longevity) may not follow a single tree. Indeed, testing for phylogenetic signal for lifespan on the CASTER topology using Pagel’s *λ* (Pagel 1999) found no significant signal (*λ* = 0.62; likelihood ratio test = 1.25, *P* = 0.26). This is to be expected for complex traits evolving on trees with high discordance (Mendes et al. 2018).

**Figure 3.**
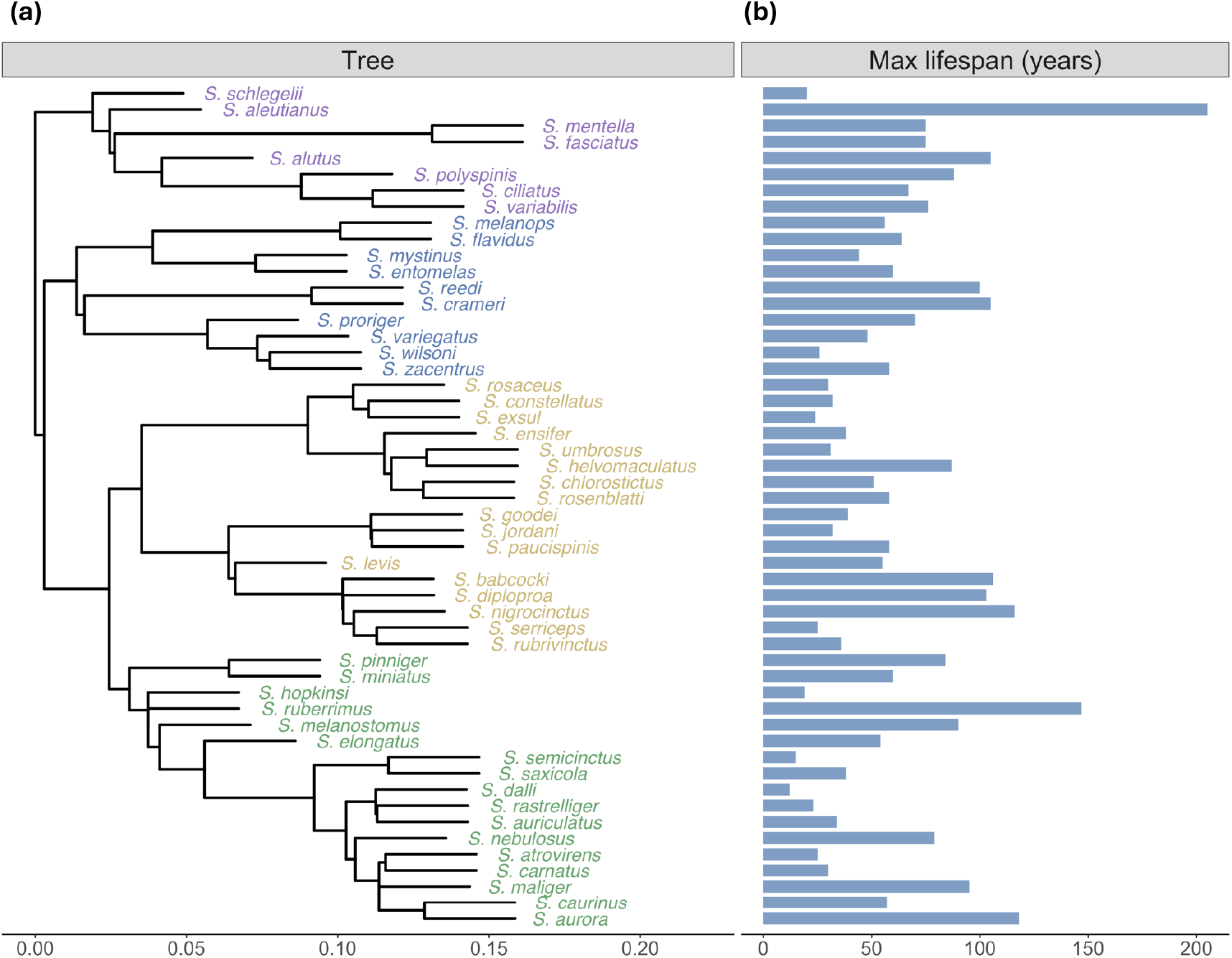
Maximum lifespan and genetic relatedness across *Sebastes*. (a) CASTER species tree in coalescent units; terminal branches were set at 0.03 coalescent units. Clade colors correspond to groups in Figure 1a. (b) Maximum lifespan for each species.

Therefore, in order to find genetic variation associated with longevity, we used a phylogenetic genome-wide association study (phyloGWAS) approach (Pease et al. 2016; Wu et al. 2018). PhyloGWAS attempts to identify variants associated with a trait, and it does so without assuming that a single species tree describes relationships at every site. We assembled biallelic nonsynonymous sites across 52 Sebastes with lifespan records (Figure 3). After filtering (see Materials and Methods), 33,716 sites from 957 genes could be tested for associations.

To account for phylogenetic relatedness without assuming a single tree (but still accounting for shared histories) we included a genetic relatedness matrix (GRM) in the linear mixed model (LMM) used for association tests. In particular, we used two methods: a tree-based GRM derived from the entire set of gene trees (*C*^∗^: Figure 4a; Hibbins et al. 2023) and a genotype-based GRM estimated directly from the variant matrix (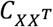 ; Figure 4b; Zhou and Stephens 2012). Both matrices recapitulate the CASTER species relationships, while also capturing relationships contributed by discordant gene trees that lack counterparts on the species tree. The tree-based GRM resolves off-diagonal structure more finely than the variant matrix-based one, reflecting the larger number of loci contributing to its estimate (Figure 4a,b).

**Figure 4.**
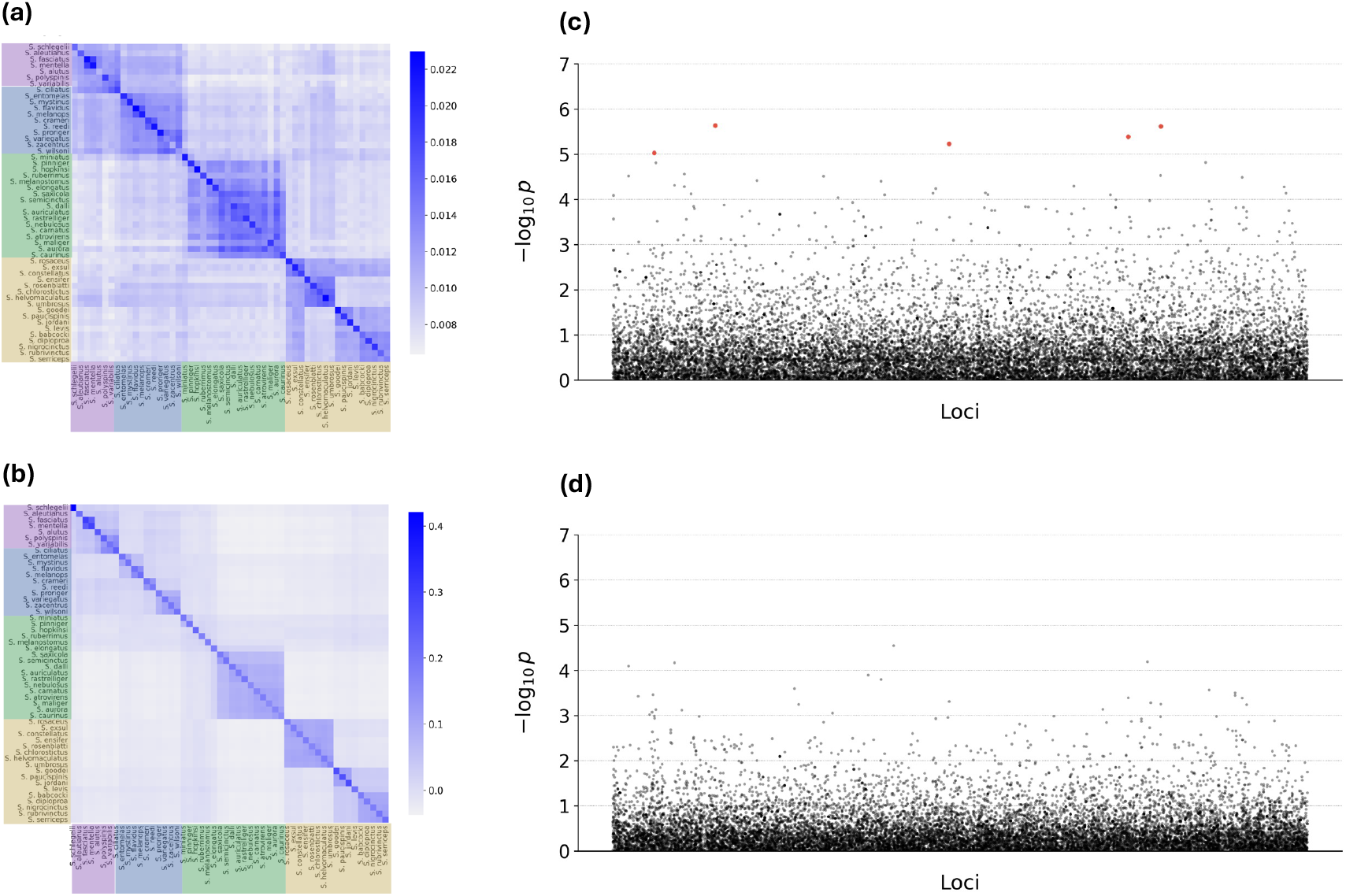
Genetic relatedness matrices and identified significant associations from the linear mixed model. Genetic relatedness matrices are inferred from (a) gene trees from 9706 loci (*C*^∗^), (b) 33,716 non-synonymous sites (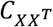). Manhattan plots show significant associations in red (*p*-values < 1×10^−5^), inferred by (c) *C*^∗^ and (d) 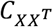. No significant associations were found in (d). The Pearson correlation coefficient is 0.69 between *p*-values in (c) and (d).

Before applying these approaches to data, we carried out simulations to examine the performance of phyloGWAS on quantitative traits; previous studies had only used discrete traits (Pease et al. 2016; Wu et al. 2018). We compared the two GRMs described above, as well as two other conditions: either using the species tree alone (*C*), or using tests that do not account for any structure—a linear model with no GRM. First, we examined the false positive rate (FPR) in our simulations (Supplementary Figure 3a). At low discordance, the linear model without a GRM showed inflated FPRs because it does not account for shared history at all. Adding any GRM reduced this inflation by about 41%. At moderate discordance, models using *C*— or 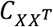 achieved the lowest FPRs, slightly better than that of *C* . At the highest discordance, all four models had very similar FPRs. Second, we examined the true positive rate (=statistical power) in our simulations. Power patterns were broadly similar across models (Supplementary Figure 3b). At a nominal *α* = 0.05, power reached up to 0.40, but declined as *α* decreased. Surprisingly, increasing discordance further reduced power for all models, and differences in power among the four models were minimal at each discordance level.

Applying linear mixed models to the rockfish data provided some promising results. Using the tree-based GRM, *C*—, we identified 5 significant nonsynonymous sites, but using the genotype-based GRM, 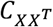, we identified 0 sites (Figure 4c,d; Supplementary Table 1). However, we found a strong positive correlation between *p*-values from both GRMs (Pearson correlation coefficient, *r*=0.69). In comparison, a linear model with no control for phylogenetic relatedness identified a very similar number (and identity) of associations as *C*— (Supplementary Figure 4; Supplementary Table 1). At the *p*-value threshold used here (=1×10^−5^), these results imply a false discovery rate of 6.7% for the results using *C*—. The sites we identified map to genes with broadly defined functions, including regulating endoplasmic reticulum stress, organizing collagen fibrils in the extracellular matrix, cell adhesion and intracellular signaling, Golgi–lysosome trafficking, and ADP ribosylation (Supplementary Table 1).

## Discussion

Genealogical discordance has become the norm in phylogenetic studies, with evidence for biological discordance due to both ILS and introgression in many different systems at many different times in the past (e.g. Pollard et al. 2006; White et al. 2009; Copetti et al. 2017; Rouard et al. 2018; Hime et al. 2021; Morales-Briones et al. 2021). Here, we found extensive genealogical discordance in *Sebastes* rockfishes. While it is quite difficult to compare the amount of discordance among systems with a single measure, as in several other published clades (e.g. Jarvis et al. 2014; Pease et al. 2016; Sun et al. 2021; Larson et al. 2025) there is no gene tree inferred here that matches the inferred species tree (any of them) at every node.

Despite extreme discordance, we expected to be able to infer a single, well-supported species tree—these are not mutually exclusive possibilities. However, none of the three methods used here agreed completely, with ∼25% of all branches differing among methods (Figure 1). It is not immediately clear to us whether there is a systematic reason the methods differ, nor which one represents (or comes closest to representing) the “true” species tree. All three methods are intended to avoid the problems of ILS, and in the limit of infinite data do avoid these problems (Mirarab et al. 2014; Chifman and Kubatko 2014; Zhang et al. 2025); ASTRAL should also not be affected by the presence of pseudoorthologous genes (Smith and Hahn 2022; Yan et al. 2022). Two of the methods, SVDquartets and CASTER, combine nucleotide sites, while the third method, ASTRAL, combines individual gene trees: this underlying difference in the data structure does not seem to lead to a clear distinction in outcomes. While many of the differences between methods represent rearrangements around just a single internal branch—and thus could be considered more minor—some rearrangements are much more severe (Supplementary Figure 1). For the minor rearrangements, it may be the internal branches with little support one way or another are simply resolved randomly by each method, as none will output a polytomy. These observed differences may then represent these random outcomes.

Although many ILS-aware methods have theoretical guarantees if discordance is due only to ILS, there are no such guarantees if introgression is also acting. Indeed, even minimal amounts of introgression can change which topology is most frequent, especially when internal branch lengths are short (Hibbins and Hahn 2024). We have not attempted to detect introgression here due to the uncertainty associated with the species tree topology. While some approaches can take such uncertainty into account (Beckman et al. 2018; Pease 2018), these are best-suited to smaller trees with a smaller number of tips. Regardless, previous studies have reported signatures of admixture among rockfishes (Schwenke et al. 2018; Wallace et al. 2022; Sykes et al. 2025), further suggesting that introgression may have contributed to the conflict found here.

Even if there was a consensus species tree agreed upon by all methods, such a tree would still not be appropriate to use as the single topology upon which all phylogenetic comparative analyses should be done. As mentioned in the Introduction, many inferences about the number and direction of evolutionary transitions will be misled in the presence of discordance. These artifacts will affect both discrete and continuous traits, and can even lead to false signals of accelerated rates of DNA sequence evolution (Mendes and Hahn 2016). In order to detect gene- or locus-wise rate changes, especially in a rapid radiation, one should use the appropriate local gene tree topology rather than the species tree topology (e.g. Yan et al. 2023).

While discordance causes many problems for comparative methods, it also offers the opportunity to uncover the genetic basis for traits (Smith et al. 2020). This occurs because discordance decouples the shared species history from each locus history, in effect randomizing genetic backgrounds. The approach used here, phyloGWAS, takes advantage of discordance to associate genetic variation with phenotypic variation. Although it had been used previously on discrete traits (Pease et al. 2016; Wu et al. 2018), here we apply phyloGWAS to a continuous trait, longevity. We used simulations to explore the power of phyloGWAS applied to continuous traits under different levels of discordance. While this approach does not increase power to detect true associations, it greatly reduced false positives compared with a linear model without relatedness. The reduction in false positives is exactly the advantage discordance provides. For instance, in the case of no discordance at all, the true mutational change(s) would be associated with the branch of the species tree common to all species carrying the phenotype of interest, but so would all other substitutions shared by those species, whether or not they were functional. Discordance can cause disparate species (i.e. not those sharing a single common ancestor in the species tree) to share a trait, thereby helping to identify functional genetic variation.

The approach to phyloGWAS used here employs a genetic relatedness matrix (GRM) in a linear mixed model (LMM) to accommodate phylogenetic structure and discordance. While previous studies have used a GRM in some phylogenetic comparative methods (Hibbins et al. 2023), Schraiber et al. (2024) showed that they could be used in a phylogenetic context in a similar way to standard GWAS. These approaches are useful in standard GWAS when there is population stratification, or other kinds of internal population structuring (Zhou and Stephens 2012). We used two approaches to constructing a GRM, finding that *C*—captured genetic relationships more accurately than 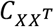, possibly because gene trees are a richer description of relationships than site patterns alone. However, 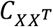 was more efficient to compute than *C*—, which becomes relevant as the number of species and gene trees increased. One disadvantage to using an LMM is that there are strong constraints on missing data. For instance, although our original data matrix had 173,332 biallelic nonsynonymous variants from 8,636 loci in *Sebastes* species with longevity phenotypes, only 33,716 sites from 957 genes could be tested for associations due to missing data. Future approaches that can better deal with the fact that genomic data will always have some missing data in some taxa are clearly needed.

We were able to find strong associations between longevity and five variants in five different genes (Supplementary Table 1). While there was no functional role shared among these genes, some are associated with tantalizing phenotypes in other systems. One variant was found in a homolog of the gene encoding Wolframin (*wfs1*), a transmembrane protein in the endoplasmic reticulum. In zebrafish models, knockout of *wfs1* leads to neurodegenerative problems associated with cell cycle delay and delayed development (Cairns et al. 2021). As increased longevity can be associated with longer development times, it is possible that *wfs1* may play a role in such changes. Similarly, we found a nonsynonymous variant in *git2a* associated with longevity. In rats, the homolog of *git2a* helps to coordinate hormonal control of aging (Chadwick et al. 2012).

Species radiations represent powerful biological laboratories for the study of speciation, adaptation, and the genotype-phenotype relationship. Here, we have taken advantage of a radiation in rockfishes to explore the genetic basis for longevity. Although there is no single clear history of speciation among these species, the genealogical histories at many thousands of loci together provide a glimpse into rapid speciation, the sorting of ancestral polymorphisms, and extreme discordance. While the our phyloGWAS approach was limited to coding regions, the further sequencing of high-quality rockfish genomes from many species will allow for greater resolution of the genotypic basis for these long-lived fishes.

## Materials and Methods

### Data sources and alignment processing

We obtained 9706 orthologous multiple-sequence alignments (MSA) covering 55 *Sebastes* species from Kolora et al. (2021). These represent a high-quality subset of all genes and species from that paper. *Sebastolobus alascanus* was added as a high-quality outgroup for 4694 loci, but no data were available for the remaining loci. Outgroup sequences were realigned to each MSA with MAFFT v7.505 using --keeplength (Katoh and Frith 2012).

### Species-tree inference and discordance metrics

Maximum-likelihood gene trees were inferred for each single-gene MSA with IQ-TREE2 (v2.3.1). Species trees were inferred with three methods: CASTER-site (v1.16.3.4; Zhang et al. 2025), ASTRAL (v1.16.3.4; Mirarab et al. 2014), and SVDquartets (implemented in PAUP* 4a168; Chifman and Kubatko 2014). Discordance was measured with site- and gene-concordance factors using IQ-TREE2 (v2.3.1), computed separately with each species tree as the fixed topology. Tree visualizations were generated with R packages phangorn v2.12.1 (Schliep 2011), phytools v2.5.2 (Revell 2012), and ggtree v3.6.2 (Yu et al. 2017).

### Genotype–phenotype association analyses

We used maximum lifespan as the phenotype for 52 *Sebastes species*. Three species (*S. minor, S. inermis*, and *S. steindachneri*) were excluded due to missing phenotypes. For genotypes, coding sequences were filtered to retain loci whose lengths were divisible by three and with zero stop codons per sequence. Filtered nucleotide MSAs were translated into amino acid alignments. Biallelic nonsynonymous sites were retained and encoded in BIMBAM format: major allele = 0, minor allele = 2, and missing genotypes = “NA”. Variants were selected with per-site missingness of less than 10% and a minor allele frequency greater than 5%.

To account for phylogenetic relationships in phyloGWAS, two GRMs were constructed: (i) A tree-based variance–covariance matrix, *C*\*, was derived from gene trees represented in substitution units across 9706 loci (Hibbins et al., 2023). (ii) A genotype-based GRM, 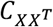, directly from the variant matrix inferred above (Zhou and Stephens 2012). Both GRMs were computed on the same set of 52 species.

Association tests were performed with linear mixed models in GEMMA (Zhou and Stephens 2012) using each GRM in turn. A linear model (no GRM) was also fit for comparison. *P*-values from GEMMA likelihood-ratio tests were evaluated at a threshold of 1×10^−5^ across sites.

### PhyloGWAS simulation settings

We carried out simulations to evaluate the performance of phyloGWAS under different levels of discordance. First, for each replicate we generated a random species tree with 50 species under a Yule process (birth rate=1×10^−5^) using Dendropy v5.0.1 (Moreno et al. 2024). For each species tree we used msprime v1.3.3 (Baumdicker et al. 2022) to simulate gene trees while varying the effective population size to generate different levels of discordance (*N*_e_=1×10^3^, 1×10^4^, 5×10^4^, 1×105, 1×10^7^). For each *N*_e_, we simulated 1000 non-recombining loci under the standard coalescent model.

We modeled a polygenic additive trait controlled by the sampled gene trees, following a similar approach to Mendes et al. (2018). Mutations were placed on each gene tree under a biallelic mutation model. When a locus contained multiple segregating sites, we randomly retained a single segregating site and discarded the others to ensure each locus contributed only once. From the full set of simulated loci, we sampled 10 variants without replacement as causal variants. For each causal variant, we drew a mutational phenotypic effect from an exponential distribution with rate parameter *λ* = 10. All descendant species inheriting the derived allele had their trait value increased by the corresponding mutational effect. Trait contributions were summed across all causal loci for each species. Finally, we added an environmental noise term drawn from a normal distribution with mean zero and variance 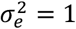 to obtain the observed trait values for each species.

To assess association methods, we compared four models: (i) a linear model (LM) with no correction for relatedness; (ii) a linear mixed model with a genetic relatedness matrix (GRM) derived from the species tree (LMM (*C*)); (iii) a linear mixed model with a GRM obtained from the collection of simulated gene trees (LMM (*C*\*)); (iv) a linear mixed model with a GRM constructed directly from the variant matrix (LMM (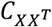)). These models allowed us to compare the impact of different choices of relatedness structure on false positive rates and true positive rates under varying levels of discordance. Association analyses were conducted with GEMMA (Zhou and Stephens 2012).

We simulated and evaluated 20 replicates for each value of *N*_e_. All scripts used for data processing, simulation, and analysis are available on GitHub at https://github.com/speechlesso/phylogwas_simulator.

## Acknowledgements

We thank Rohit Kolora, Runyang Lou, and Greg Owens for sharing data and helpful discussions. This research was funded by National Science Foundation grant DBI-2146866.

## References

Adams, R., Lozano, J. R., Duncan, M., Green, J., Assis, R., & DeGiorgio, M. (2025). A tale of too many trees: a conundrum for phylogenetic regression. Molecular Biology and Evolution, 42(3), msaf032.

Ané, C., Larget, B., Baum, D. A., Smith, S. D., & Rokas, A. (2007). Bayesian estimation of concordance among gene trees. Molecular Biology and Evolution, 24(2), 412–426.

Baumdicker, F., Bisschop, G., Goldstein, D., Gower, G., Ragsdale, A. P., Tsambos, G., … & Kelleher, J. (2022). EVicient ancestry and mutation simulation with msprime 1.0. Genetics, 220(3), iyab229.

Beckman, E. J., Benham, P. M., Cheviron, Z. A., & Witt, C. C. (2018). Detecting introgression despite phylogenetic uncertainty: the case of the South American siskins. Molecular Ecology, 27(22), 4350–4367.

Cairns, G., Burté, F., Price, R., O’Connor, E., Toms, M., Mishra, R., … & Yu-Wai-Man, P. (2021). A mutant wfs1 zebrafish model of Wolfram syndrome manifesting visual dysfunction and developmental delay. Scientific Reports, 11(1), 20491.

Chadwick, W., Martin, B., Chapter, M. C., Park, S. S., Wang, L., Daimon, C. M., … & Maudsley, S. (2012). GIT2 acts as a potential keystone protein in functional hypothalamic networks associated with age-related phenotypic changes in rats. PLoS ONE, 7(5), e36975.

Chifman, J., & Kubatko, L. (2014). Quartet inference from SNP data under the coalescent model. Bioinformatics, 30(23), 3317–3324.

Copetti, D., Búrquez, A., Bustamante, E., Charboneau, J. L., Childs, K. L., Eguiarte, L. E., … & Sanderson, M. J. (2017). Extensive gene tree discordance and hemiplasy shaped the genomes of North American columnar cacti. Proceedings of the National Academy of Sciences, 114(45), 12003–12008.

Degnan, J. H., & Rosenberg, N. A. (2009). Gene tree discordance, phylogenetic inference and the multispecies coalescent. Trends in Ecology & Evolution, 24(6), 332–340.

Doyle, J. J. (1992). Gene trees and species trees: molecular systematics as one-character taxonomy. Systematic Botany, 144–163.

Felsenstein, J. (1985). Phylogenies and the comparative method. The American Naturalist, 125(1), 1–15.

Feng, S., Bai, M., Rivas-González, I., Li, C., Liu, S., Tong, Y., … & Zhang, G. (2022). Incomplete lineage sorting and phenotypic evolution in marsupials. Cell, 185(10), 1646–1660.

Fontaine, M. C., Pease, J. B., Steele, A., Waterhouse, R. M., Neafsey, D. E., Sharakhov, I. V., … & Besansky, N. J. (2015). Extensive introgression in a malaria vector species complex revealed by phylogenomics. Science, 347(6217), 1258524.

Gomez-Uchida, D., & Banks, M. A. (2006). Estimation of eVective population size for the long-lived darkblotched rockfish Sebastes crameri. Journal of Heredity, 97(6), 603–606.

Guerrero, R. F., & Hahn, M. W. (2018). Quantifying the risk of hemiplasy in phylogenetic inference. Proceedings of the National Academy of Sciences, 115(50), 12787–12792.

Hahn, M. W., & Nakhleh, L. (2016). Irrational exuberance for resolved species trees. Evolution, 70(1), 7–17.

Han, F., Lamichhaney, S., Grant, B. R., Grant, P. R., Andersson, L., & Webster, M. T. (2017). Gene flow, ancient polymorphism, and ecological adaptation shape the genomic landscape of divergence among Darwin’s finches. Genome Research, 27(6), 1004–1015.

Hibbins, M. S., Gibson, M. J., & Hahn, M. W. (2020). Determining the probability of hemiplasy in the presence of incomplete lineage sorting and introgression. eLife, 9, e63753.

Hibbins, M. S., Breithaupt, L. C., & Hahn, M. W. (2023). Phylogenomic comparative methods: accurate evolutionary inferences in the presence of gene tree discordance. Proceedings of the National Academy of Sciences, 120(22), e2220389120.

Hibbins, M. S., & Hahn, M. W. (2024). Distinguishing between histories of speciation and introgression using genomic data, Bulletin of the Society of Systematic Biologists 3(1), doi: 10.18061/bssb.v3i1.9227.

Hime, P. M., Lemmon, A. R., Lemmon, E. C. M., Prendini, E., Brown, J. M., Thomson, R. C., … & Weisrock, D. W. (2021). Phylogenomics reveals ancient gene tree discordance in the amphibian tree of life. Systematic Biology, 70(1), 49–66.

Hyde, J. R., & Vetter, R. D. (2007). The origin, evolution, and diversification of rockfishes of the genus Sebastes (Cuvier). Molecular Phylogenetics and Evolution, 44(2), 790–811.

Jarvis, E. D., Mirarab, S., Aberer, A. J., Li, B., Houde, P., Li, C., … & Zhang, G. (2014). Whole-genome analyses resolve early branches in the tree of life of modern birds. Science, 346(6215), 1320–1331.

Johns, G. C., & Avise, J. C. (1998). Tests for ancient species flocks based on molecular phylogenetic appraisals of Sebastes rockfishes and other marine fishes. Evolution, 52(4), 1135–1146.

Katoh, K., & Frith, M. C. (2012). Adding unaligned sequences into an existing alignment using MAFFT and LAST. Bioinformatics, 28(23), 3144–3146.

Kolora, S. R. R., Owens, G. L., Vazquez, J. M., Stubbs, A., Chatla, K., Jainese, C., … & Sudmant, P. H. (2021). Origins and evolution of extreme life span in Pacific Ocean rockfishes. Science, 374(6569), 842–847.

Lamichhaney, S., Han, F., Berglund, J., Wang, C., Almén, M. S., Webster, M. T., … & Andersson, L. (2016). A beak size locus in Darwin’s finches facilitated character displacement during a drought. Science, 352(6284), 470–474.

Lanfear, R., & Hahn, M. W. (2024). The meaning and measure of concordance factors in phylogenomics. Molecular Biology and Evolution, 41(11), msae214.

Larson, D. A., Staton, M. E., Kapoor, B., Islam-Faridi, N., Zhebentyayeva, T., Fan, S., … & Nelson, C. D. (2025). A haplotype-resolved reference genome of Quercus alba sheds light on the evolutionary history of oaks. New Phytologist, 246(1), 331–348.

Li, Y. Y., Liu, Z., Qi, F. Y., Zhou, X., & Shi, P. (2016). Functional eVects of a retained ancestral polymorphism in Prestin. Molecular Biology and Evolution, 34(1), 88–92.

Louis, M., Galimberti, M., Archer, F., Berrow, S., Brownlow, A., Fallon, R., … & Gaggiotti, O. E. (2021). Selection on ancestral genetic variation fuels repeated ecotype formation in bottlenose dolphins. Science Advances, 7(44), eabg1245.

Love, M. S., Yoklavich, M., & Thorsteinson, L. K. (2002). The rockfishes of the northeast Pacific. Univ of California Press.

Maddison, W. P. (1997). Gene trees in species trees. Systematic Biology, 46(3), 523–536.

Mangel, M., Kindsvater, H. K., & Bonsall, M. B. (2007). Evolutionary analysis of life span, competition, and adaptive radiation, motivated by the Pacific rockfishes (Sebastes). Evolution, 61(5), 1208–1224.

Mendes, F. K., Fuentes-González, J. A., Schraiber, J. G., & Hahn, M. W. (2018). A multispecies coalescent model for quantitative traits. eLife, 7, e36482.

Minh, B. Q., Schmidt, H. A., Chernomor, O., Schrempf, D., Woodhams, M. D., Von Haeseler, A., & Lanfear, R. (2020a). IQ-TREE 2: new models and eVicient methods for phylogenetic inference in the genomic era. Molecular Biology and Evolution, 37(5), 1530–1534.

Minh, B. Q., Hahn, M. W., & Lanfear, R. (2020b). New methods to calculate concordance factors for phylogenomic datasets. Molecular Biology and Evolution, 37(9), 2727–2733.

Mirarab, S., Reaz, R., Bayzid, M. S., Zimmermann, T., Swenson, M. S., & Warnow, T. (2014). ASTRAL: genome-scale coalescent-based species tree estimation. Bioinformatics, 30(17), i541–i548.

Mo, Y. K., Lanfear, R., Hahn, M. W., & Minh, B. Q. (2023). Updated site concordance factors minimize eVects of homoplasy and taxon sampling. Bioinformatics, 39(1), btac741.

Mo, Y. K., & Hahn, M. W. (2026). Estimating the rate of quantitative trait evolution in the presence of gene tree discordance by calculating likelihoods across trees. Evolutionary Journal of the Linnean Society, kzag004

Morales-Briones, D. F., Kadereit, G., Tefarikis, D. T., Moore, M. J., Smith, S. A., Brockington, S. F., … & Yang, Y. (2021). Disentangling sources of gene tree discordance in phylogenomic data sets: testing ancient hybridizations in Amaranthaceae s.l. Systematic Biology, 70(2), 219–235.

Moreno, M. A., Holder, M. T., & Sukumaran, J. (2024). DendroPy 5: a mature Python library for phylogenetic computing. Journal of Open Source Software, 9(101), 6943

Pagel, M. (1999). Inferring the historical patterns of biological evolution. Nature, 401(6756), 877–884.

Palesch, D., Bosinger, S. E., Tharp, G. K., Vanderford, T. H., Paiardini, M., Chahroudi, A., … & Silvestri, G. (2018). Sooty mangabey genome sequence provides insight into AIDS resistance in a natural SIV host. Nature, 553(7686), 77–81.

Pease, J. B., Haak, D. C., Hahn, M. W., & Moyle, L. C. (2016). Phylogenomics reveals three sources of adaptive variation during a rapid radiation. PLoS Biology, 14(2), e1002379.

Pease, J. B. (2018). Why phylogenomic uncertainty enhances introgression analyses. Molecular Ecology, 27(22), 4347–4349.

Pollard, D. A., Iyer, V. N., Moses, A. M., & Eisen, M. B. (2006). Widespread discordance of gene trees with species tree in Drosophila: evidence for incomplete lineage sorting. PLoS Genetics, 2(10), e173.

Revell, L. J. (2012). phytools: an R package for phylogenetic comparative biology (and other things). Methods in Ecology and Evolution, 3(2), 217–223.

Rouard, M., Droc, G., Martin, G., Sardos, J., Hueber, Y., Guignon, V., … & Roux, N. (2018). Three new genome assemblies support a rapid radiation in Musa acuminata (wild banana). Genome Biology and Evolution, 10(12), 3129–3140.

Schliep, K. P. (2011). phangorn: phylogenetic analysis in R. Bioinformatics, 27(4), 592–593.

Schraiber, J. G., Edge, M. D., & Pennell, M. (2024). Unifying approaches from statistical genetics and phylogenetics for mapping phenotypes in structured populations. PLoS Biology, 22(10), e3002847.

Schwenke, P. L., Park, L. K., & Hauser, L. (2018). Introgression among three rockfish species (Sebastes spp.) in the Salish Sea, northeast Pacific Ocean. PloS ONE, 13(3), e0194068.

Smith, S. D., Pennell, M. W., Dunn, C. W., & Edwards, S. V. (2020). Phylogenetics is the new genetics (for most of biodiversity). Trends in Ecology and Evolution, 35(5), 415–425.

Smith, M. L., & Hahn, M. W. (2022). The frequency and topology of pseudoorthologs. Systematic Biology, 71(3), 649–659.

Sun, C., Huang, J., Wang, Y., Zhao, X., Su, L., Thomas, G. W., … & Mueller, R. L. (2021). Genus-wide characterization of bumblebee genomes provides insights into their evolution and variation in ecological and behavioral traits. Molecular Biology and Evolution, 38(2), 486–501.

Sykes, N. T., Lou, R. N., Siegle, M. R., Sudmant, P. H., Larson, W. A., & Owens, G. L. (2025). Phylogenetically diverse introgression drives subtle population structure in Pacific rockfishes. bioRxiv, doi: 10.1101/2025.09.23.678143.

Treaster, S., Deelen, J., Daane, J. M., Murabito, J., Karasik, D., & Harris, M. P. (2023). Convergent genomics of longevity in rockfishes highlights the genetics of human life span variation. Science Advances, 9(2), eadd2743.

Wallace, E. N., Reed, E. M. X., Aguilar, A., & Reiskind, M. B. (2022). Resolving the phylogenetic relationship among recently diverged members of the rockfish subgenus Sebastosomus. Molecular Phylogenetics and Evolution, 173, 107515.

Waterhouse, R. M., Seppey, M., Simão, F. A., Manni, M., Ioannidis, P., Klioutchnikov, G., … & Zdobnov, E. M. (2018). BUSCO applications from quality assessments to gene prediction and phylogenomics. Molecular Biology and Evolution, 35(3), 543–548.

White, M. A., Ané, C., Dewey, C. N., Larget, B. R., & Payseur, B. A. (2009). Fine-scale phylogenetic discordance across the house mouse genome. PLoS Genetics, 5(11), e1000729.

Wu, M., Kostyun, J. L., Hahn, M. W., & Moyle, L. C. (2018). Dissecting the basis of novel trait evolution in a radiation with widespread phylogenetic discordance. Molecular Ecology, 27(16), 3301–3316.

Yan, Z., Smith, M. L., Du, P., Hahn, M. W., & Nakhleh, L. (2022). Species tree inference methods intended to deal with incomplete lineage sorting are robust to the presence of paralogs. Systematic Biology, 71(2), 367–381.

Yan, H., Hu, Z., Thomas, G. W., Edwards, S. V., Sackton, T. B., & Liu, J. S. (2023). PhyloAcc-GT: a Bayesian method for inferring patterns of substitution rate shifts on targeted lineages accounting for gene tree discordance. Molecular Biology and Evolution, 40(9), msad195.

Yu, G., Smith, D. K., Zhu, H., Guan, Y., & Lam, T. T. Y. (2017). ggtree: an R package for visualization and annotation of phylogenetic trees with their covariates and other associated data. Methods in Ecology and Evolution, 8(1), 28–36.

Zhang, C., Nielsen, R., & Mirarab, S. (2025). CASTER: Direct species tree inference from whole-genome alignments. Science, 387(6737), eadk9688.

Zhou, X., & Stephens, M. (2012). Genome-wide eVicient mixed-model analysis for association studies. Nature Genetics, 44(7), 821–824.

