## Supplementary Figures and Tables for "Linking genotype to longevity under genealogical discordance in *Sebastes* rockfishes"

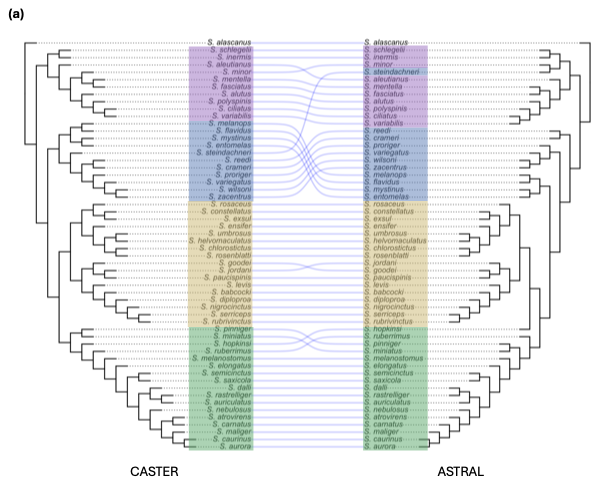

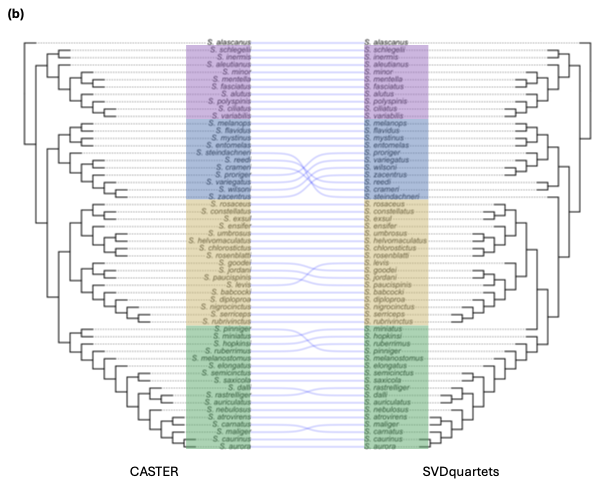

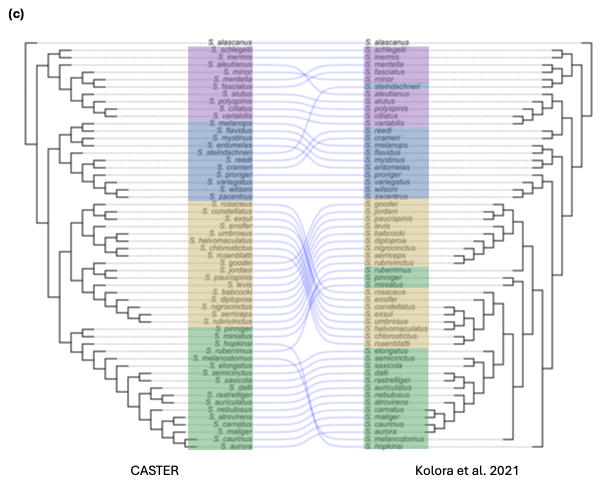

Supplementary Figure 1. Tanglegrams comparing species trees from alternative methods. Across panels, the CASTER topology (left) is fixed; clade colors follow the CASTER clades. All branch lengths are unit length.

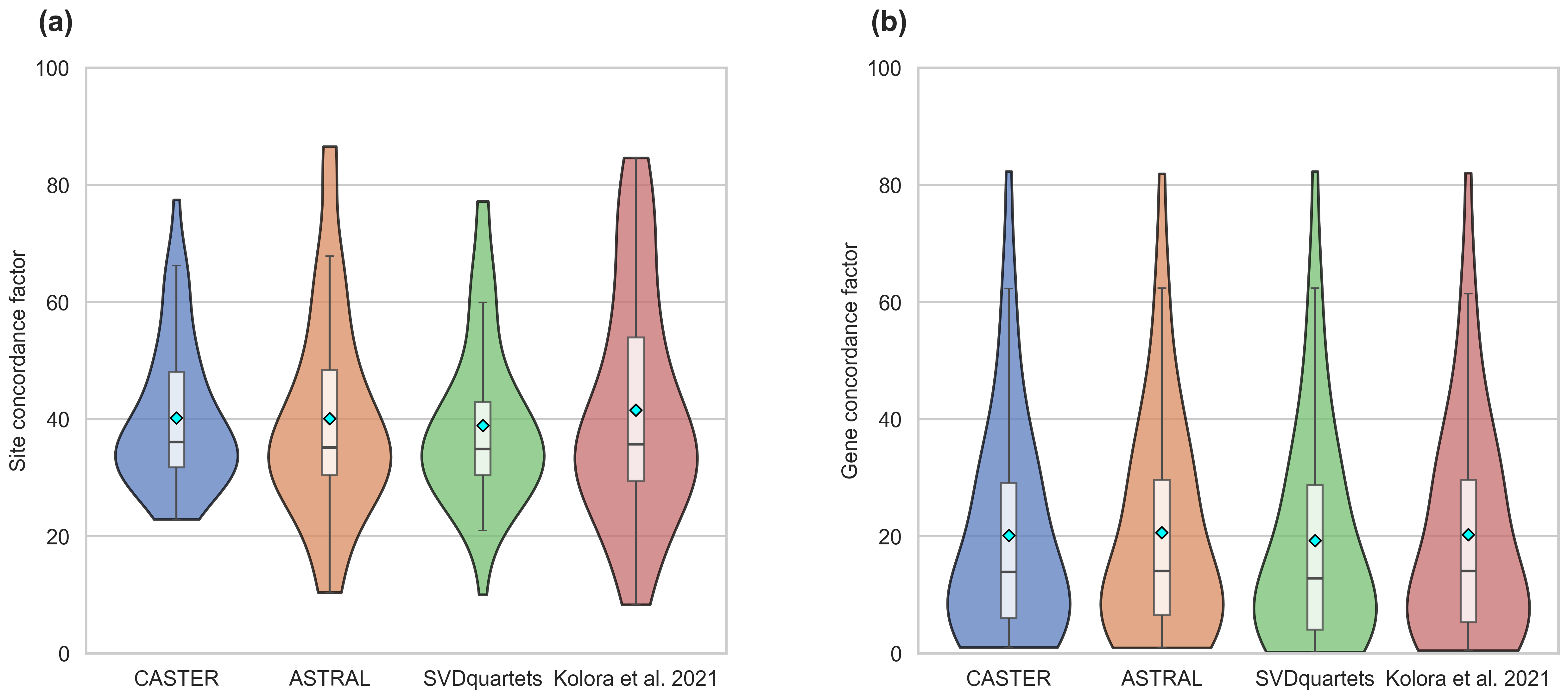

Supplementary Figure 2. Distributions of concordance factors for (a) site concordance factors and (b) gene concordance factors, across species trees inferred by CASTER, ASTRAL, SVDquartets, and Kolora et al. (2021). Diamonds denote means.

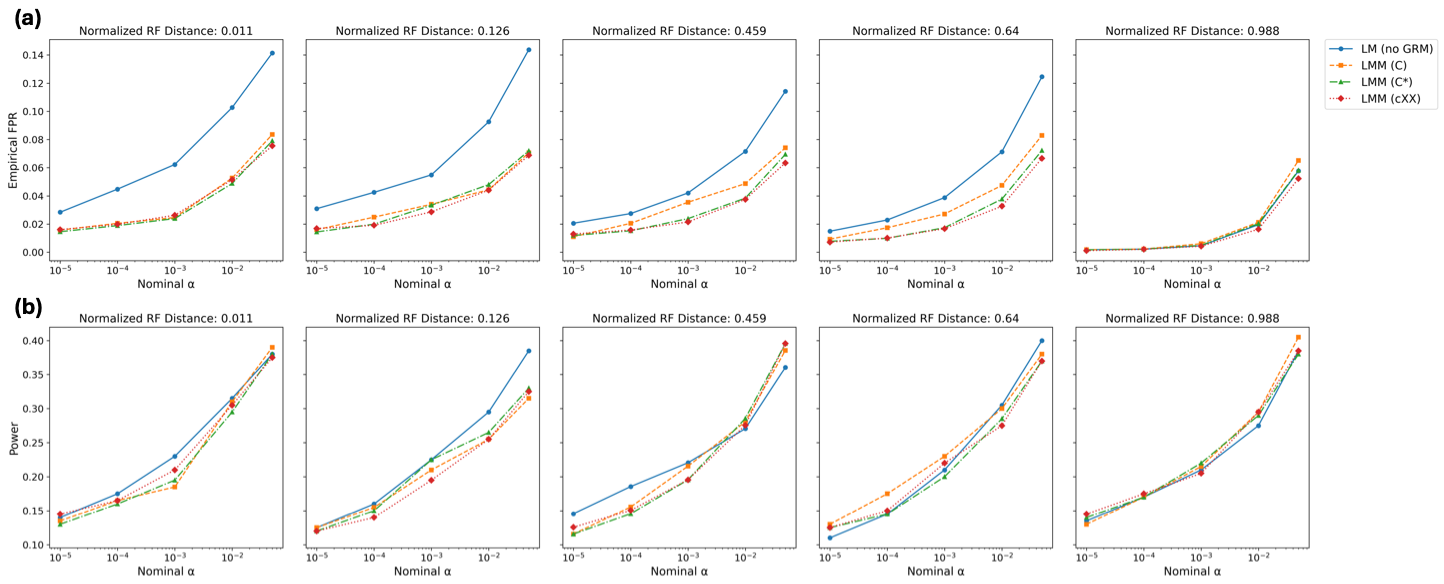

Supplementary Figure 3. Performance of phyloGWAS in simulations under varying genealogical discordance. We examined (a) empirical false positive rate (FPR) and (b) true positive rate (power) across different *p*-value cutoffs (nominal α) for four models: linear model without a GRM (LM), and linear mixed models with $C$, $C^{*}$, or $C_{XX^{T}}$ as the GRM. From left to right, panels represent simulations with increasing discordance; this discordance is summarized as the normalized Robinson-Foulds distance of gene trees to the species tree in each case.

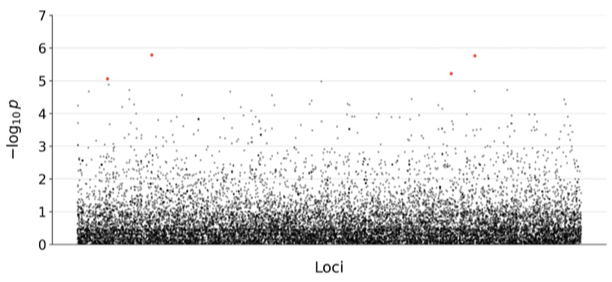

Supplementary Figure 4: Identified significant associated sites from the linear model, without a genetic relatedness matrix.

Supplementary Table 1: Identified longevity-associated nonsynonymous variants.

| Ortholog Group | Variant | LMM ($C^{*}$)^a^ | LM ^a^ | Gene | Description |
| --- | --- | --- | --- | --- | --- |
| 4579 | K/R | ✓ | ✓ | wfs1b | Wolfram syndrome 1b (wolframin) |
| 14501 | F/Y | ✓ | ✓ | LOC115024838 | decorin-like |
| 13862 | G/S | ✓ | ✓ | git2a | G protein-coupled receptor kinase interacting ArfGAP 2a |
| 9988 | T/Q | ✓ | 🗶 | gaa2 | alpha glucosidase 2 |
| 3178 | A/P | ✓ | ✓ | LOC113145053 | protein mono-ADP-ribosyltransferase PARP14-like |

^a^A checkmark indicates that the variant was significant using this method
